# Tadpole-like conformations of huntingtin exon 1 with expanded polyglutamine engenders novel interactions in cells

**DOI:** 10.1101/179663

**Authors:** Estella A. Newcombe, Kiersten M. Ruff, Ashish Sethi, Angelique R. Ormsby, Yasmin M. Ramdzan, Archa Fox, Anthony W. Purcell, Paul R. Gooley, Rohit V. Pappu, Danny M. Hatters

## Abstract

Soluble huntingtin exon 1 (Httex1) with expanded polyglutamine (polyQ) engenders neurotoxicity in Huntington’s disease. To uncover the physical basis of this toxicity, we performed structural studies of soluble Httex1 for wild type and mutant polyQ lengths. Nuclear magnetic resonance experiments show evidence for conformational rigidity across the polyQ region. In contrast, hydrogen-deuterium exchange shows absence of backbone amide protection, suggesting negligible persistence of hydrogen bonds. The seemingly conflicting results are explained by all-atom simulations, which show that Httex1 adopts *tadpole-like* structures with a globular head encompassing the N-terminal amphipathic and polyQ regions and the tail encompassing the C-terminal proline-rich region. The surface area of the globular domain increases monotonically with polyQ length. This stimulates sharp increases in gain-of-function interactions in cells for expanded polyQ, and one of these interactions is with the stress-granule protein Fus. Our results highlight plausible connections between Httex1 structure and routes to neurotoxicity.

---

Huntington’s Disease (HD) is caused by mutations in exon 1 of the huntingtin (Htt) gene that expand a CAG trinucleotide repeat sequence, encoding polyglutamine (polyQ), from a normal range of 11 – 25 to lengths beyond 36 (1). A key feature of HD is the appearance of intracellular aggregates of N-terminal fragments of mutant Htt that includes the polyQ segment (2). Ectopic expression of exon 1 in its polyQ-expanded form is sufficient to recapitulate neurological defects and aggregation pathology resembling HD in a variety of animal model contexts (3-5). Because of this, the Htt exon 1 fragment (Httex1), has been the focus of intensive study in efforts to illuminate the basis of the “gain-of-toxicity” functions attributable to the polyQ expansion.

There is growing evidence that toxic properties of mutant Htt arise from the interactions involving soluble forms of polyQ-expanded Httex1 prior to its aggregation into visible aggregates (6-13). Yet, an outstanding question remains: how do soluble forms of Httex1 that encompass expanded polyQ tracts give rise to toxicity and cell death? Some of the most popular hypotheses stem from studies using monoclonal antibodies to measure distinct conformational epitopes of Httex1 (9, 10, 12, 14). These studies point to polyQ expansion enabling access to novel conformations that mediate pathogenic interactions. However, other studies have argued that binding to expanded polyQ constructs results from expanded polyQ constructs having an increased number of binding sites compared to wild-type polyQ constructs (15, 16). Overall, the resolution of antibody-based studies is inadequate to make precise assessments of the impact of polyQ length on the conformational properties of monomeric Httex1.

Clearly, additional detailed investigations of soluble forms of Httex1 are required to guide structure-function studies that connect features within soluble forms of Httex1 to cellular toxicity. Such studies are challenging because of the sequence-encoded conformational heterogeneity of Httex1 (17-19), the repetitive nature of amino acid sequence blocks within Httex1, and the high aggregation propensity of molecules with polyQ stretches (20). To overcome problems with aggregation, many researchers have resorted to studying constructs housing short polyQ tracts or using fusion proteins wherein solubilizing folded domains are fused to Httex1 constructs or polyQ peptides of different lengths (21-23). None of these studies have provided clear insights regarding the connection between expanded polyQ and the toxic attributes to the soluble Httex1 protein. For example, one-dimensional nuclear magnetic resonance (NMR) experiments on short (22Q) versus long (41Q) polyQ sequences as fusions to Glutathione S-transferase (GST) yielded no apparent differences in solution structures (24). Crystals of Httex1 (17Q) fused to maltose binding protein showed the presence of multiple structures indicating that polyQ adopts different conformations, including α-helical structures that extend from the N-terminal end (25). Yet this study did not report structures of pathogenic polyQ lengths.

Our goal is to obtain a structural description of the biologically relevant sequence of Httex1, without fusion domains, and to understand how polyQ expansions alter the structural features of soluble forms of this molecule. For this, we used a combination of hydrogen-deuterium exchange (HDX) with NMR and mass spectrometry and studied two variants *viz*., 25Q and 46Q of Httex1. Regardless of polyQ length, Httex1 showed a lack of persistent secondary structure, yet the N17 and polyQ domains displayed unusual rigidity. By complementing these solution-phase studies with atomistic simulations we found that monomeric Httex1 forms *tadpole-like* structures involving a globular head, characterized by the adsorption of the amphipathic 17-residue N-terminal sequence (N17) on a globular polyQ domain, and a semi-flexible proline-rich C-terminal tail. This tadpole-like topology is preserved for wild type and expanded polyQ lengths. However, the overall size and surface area of the globular polyQ domain grows monotonically with the length of polyQ tracts. Accordingly, we hypothesized that the growing prominence of the polyQ domain in tadpole-like topologies could engender gain-of-function interactions between soluble Httex1 and components of the cellular proteome. We tested this hypothesis using a proteomic analysis and found that Httex1 with longer polyQ sequences leads to gain-of-interactions with other proteins. Notable among the gain-of-function interactions are those with the RNA binding protein Fus and other RNA binding proteins. We showed that the gain-of-function interactions with Fus does become protective inasmuch as overexpression of Fus increases cell survival by reducing the toxicity associated with soluble forms of Httex1. The biophysical model emerging from our work suggests that the tadpole-like architecture for Httex1 and the monotonically increasing surface area (and number of binding sites) of the polyQ domain promotes increased gain-of-function interactions involving the soluble forms of Httex1. These interactions can become protective, as demonstrated here with the overexpression of Fus, or toxic, which is the more likely scenario for endogenous levels of expression of proteins in the gain-of-function network. This toxicity would derive from the recruitment and sequestration of essential proteins, including a variety of transcriptional regulators (26-28).

## RESULTS

### PolyQ expansion does not alter Httex1 structural (dis)order

Sequence-based predictors suggest that Httex1 is intrinsically disordered – a prediction that is insensitive to increases in polyQ length (Fig S1) (29). If the predictors are correct, then Httex1 should generally lack stable hydrogen bonds. Hydrogen-deuterium exchange (HDX), which when combined with NMR spectroscopy or mass spectrometry (MS) serves as a powerful method to detect the presence of stable hydrogen bonds. We used HDX combined with NMR to uncover protected amide hydrogens for the 25Q construct. Utilizing purified recombinant Httex1 25Q (Fig S2; Table S1) we followed HDX upon dissolving protonated Httex1 in 100% deuterated buffer and probed for loss of NH resonances by ^1^H-^15^N HSQC. At pH 7.5, no resonances were protected from exchange for either the 25Q or 46Q forms, suggesting an absence of persistent hydrogen bonds (Fig S3). In an effort to detect weak hydrogen bonds we also investigated HDX at pH 4. Under these conditions, the exchange of a freely exposed peptide group is approximately 3-orders of magnitude slower than at pH 7.5 (30). For Httex1 25Q, seven backbone amide hydrogens were protected from deuteration at pH 4. These correspond to residues in the C-terminal region L67, L68, Q72, G83, L91, V86, and W97 (Fig 1A). Except for the C-terminal W97 all these residues with protected amide hydrogens were flanked on one or both sides by proline residues. In addition, the side-chain NH_2_ groups of Gln residues also exchanged rapidly.

**Figure 1.**
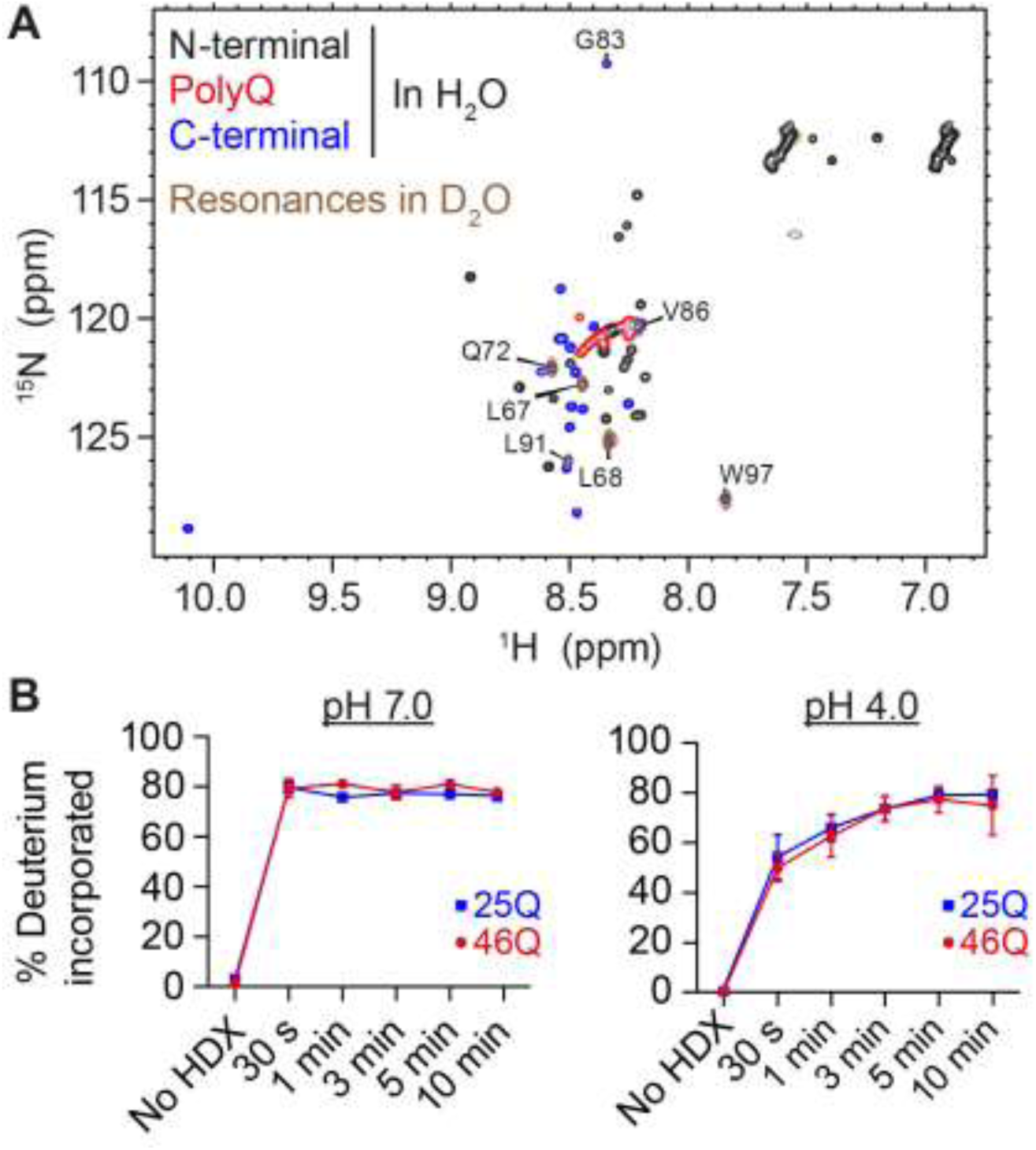
Httex1 lacks persistent hydrogen bonding with 25Q or 46Q. (**A**) ^1^H,^15^N HSQC of Httex1 protein fragment in H_2_O versus D_2_O based sodium acetate buffer (150 mM; pH 4). Labelled residues are observed after hydrogen-deuterium exchange. (**B**) Hydrogen-deuterium exchange of Httex1 25Q and 46Q were measured over 10 minutes using mass spectroscopy at both pH 7.5 and pH 4. Exchange plateaued at ∼80% (due to back-exchange in the protonated solvent during chromatography separation prior to MS).

We could not obtain an NMR spectrum for the 46Q form of Httex1 because of two technical challenges. First, the protein aggregated at the high concentrations (100 μM) required for NMR. Second, it was not possible to assign every Gln in the polyQ stretch to a unique resonance (Fig 2A). As an alternative strategy to measure structural differences between the 46Q and 25Q forms of Httex1 we turned to HDX-mass spectrometry (HDX-MS; Fig S4). This approach enabled measurements of global HDX under far more dilute protein concentrations (5 μM) whereby aggregation was considerably slower and hence measurements could be attained before aggregation. There was no significant difference in HDX between the 25Q and 46Q forms whether the measurements were performed at pH 7.5 or pH 4 (Fig 1B). Treatment with denaturant (8 M urea-d_4_) also did not change the HDX patterns. These results collectively indicate that Httex1 is devoid of persistent hydrogen bonding in its soluble form and this is true irrespective of whether the polyQ length is in the normal or pathological range.

**Figure 2.**
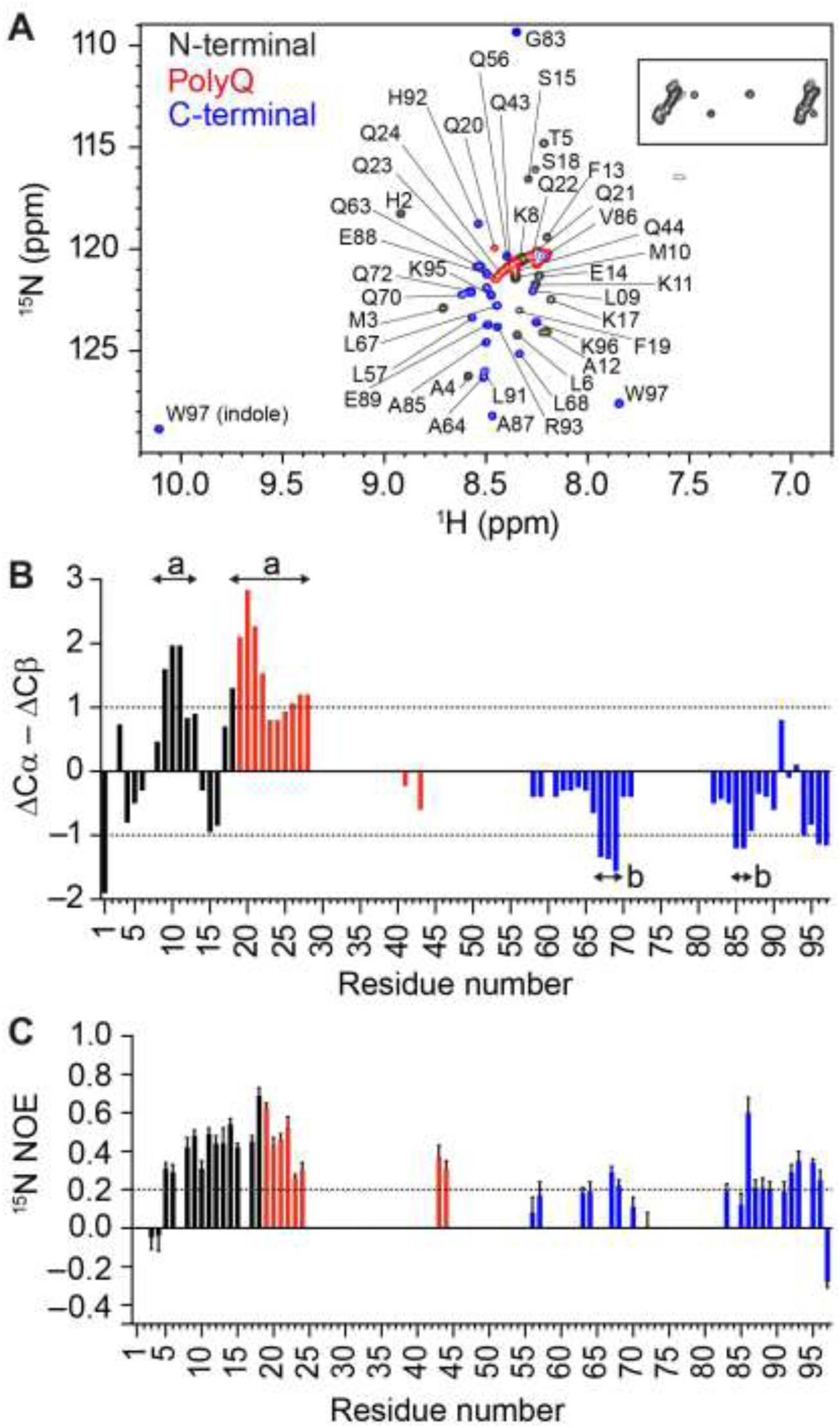
Resonance assignment of the Httex1 monomer suggests presence of transient structure. (**A**) ^1^H,^15^N HSQC of Httex1 protein fragment (25Q) in sodium acetate buffer (150 mM; pH 4), recorded at 5 °C, indicating N-terminal, polyQ and C-terminal Httex1 backbone amide peaks, with glutamine side chains (boxed). (**B**) Analysis of propensity of secondary structure based on ΔCα – ∆Cβ values where persistence of positive ∆Cα – ∆Cβ differences suggest α-helical structures and persistence of negative ∆Cα – ∆Cβ differences are consistent with β-strands. (**C**) ^15^N{^1^H}-NOE revealed positive NOE values of 0.4 to 0.6 for N-terminal Httex1 25Q and part of the polyQ region.

### Soluble polyQ is devoid of persistent secondary structure but is characterized by the presence of rigid domains

Previous experiments and simulations have suggested that polyQ prefers to adopt an array of collapsed conformations of equivalent thermodynamic preference (31, 32) that arises from water being a poor solvent for polyQ (33). Collapsed globular conformations minimize the chain-solvent interface between polyQ and aqueous solvent. This behaviour is in contrast to that of canonical random coils that prefer expanded conformations to maximize chain-solvent interactions (34). Additionally, studies based on atomic force microscopy suggest that polyQ globules display considerable mechanical rigidity (19, 35, 36). Importantly, these compact structures have been previously suggested by simulations to be formed via labile combinations of backbone and side-chain hydrogen bonds whereby stable hydrogen bonds do not persist (31, 37-39), an observation that is also true of generic polyamides including polypeptide backbones sans any side-chains (40).

Our HDX data are consistent with predictions of conformational heterogeneity within the polyQ domain, and through Httex1. To understand the broader implications of our HDX data and discern differences, if any, in the sequence-specific conformational preferences of different sequence blocks within Httex1 we used high-resolution NMR spectroscopy. Fig 2A shows the near complete non-proline backbone assignment of ^13^C,^15^N-labelled-Httex1 (25Q) using triple resonance NMR experiments (41). Approximately 90% of Httex1 sequence, excluding the polyQ region and proline residues, was assignable (Fig 2A; Table S2). Although we could not assign individual backbone resonances within the polyQ tract, the side-chain resonances were readily identifiable as overlapping resonances (between ^1^H of 6.8–7.8 ppm and ^15^N of 111.5–114 ppm; Fig 2A, boxed). Additionally, the backbone, based on ^13^Cα–β peak correlations, formed a resonance arc (between ^1^H 8.17–8.49 ppm and ^15^N 120–121.7 ppm; Fig 2A, red). The ^13^Cα and ^13^Cβ chemical shifts are sensitive to secondary structure (42). Analysis of the smoothed ΔCα–ΔCβ values (43) indicated localized regions of distinct secondary structure and regions lacking structure with a tendency towards some α-helical structure throughout the N-terminal and polyQ regions of Httex1, as indicated by values that are ≥ 1 (42-44). This result is consistent with the findings that portions of the polyQ region are able to form α-helices (25, 45, 46). In light of the lack of protection from HDX, the simplest explanation is that the α-helical segments form transiently. The remaining C-terminal sequence displayed clear preferences for extended structures (Fig 2B). Overall, these results suggest that Httex1 adopts a heterogeneous ensemble of conformations with some underlying residual structure.

To investigate the issue of molecular rigidity anticipated from the preference for globular conformations within the polyQ domain in particular, we assessed the backbone heteronuclear Overhauser effect (^15^N{^1^H}-NOE) for the 25Q construct. Positive NOE values (> 0.85) indicate a rigid protein backbone (Fig S5). The first four residues within Httex1 have negative NOE values, which is consistent with high mobility of the N-terminus (Fig 2C). The next 13 amino acids N-terminal to the polyQ region as well as the arc of glutamine backbone resonances displayed positive NOE values mostly between 0.4 and 0.6. The C-terminal proline-rich region show ^5^N{^1^H}-NOE values <0.4. These observations suggest that, while the N17 and polyQ tract are not ordered in the canonical sense, there is a persistent preference for conformational rigidity or more precisely, low amplitude conformational fluctuations, especially within the N17 and polyQ regions. In comparison, other intrinsically disordered proteins including the synucleins (47, 48) and PAGE4 (49) show ^15^N{^1^H}-NOE values of 0.1 to 0.4, similar to the proline-rich region of Httex1. Overall, our data point to an interesting duality whereby Httex1 molecules sample conformations in aqueous solutions with labile intra-molecular hydrogen bonds, although these conformations also involve a coexistence of compact-rigid parts within N17 and the polyQ tract and extended conformations within the proline-rich region. These findings hint at a tadpole-like architecture for Httex1, and we assessed this possibility using atomistic simulations.

### Atomistic simulations highlight distinctive tadpole-like architectures for Httex1

We preformed atomistic simulations of Httex1 with 25Q and 46Q using the ABSINTH implicit solvation model and force field paradigm (50) to provide a physical picture for the dualities observed using HDX versus ΔCα–ΔCβ and ^15^N{^1^H}NOE analysis. The results are summarized in Fig 3. Both 25Q and 46Q showed limited persistent secondary structural preferences (Fig 3A). A modest preference for α-helical structure extending from the N-terminus through the polyQ domain was observed, and this is consistent with the Δ Cα–ΔCβ profile for 25Q. To quantify the average architecture of Httex1, we quantified the ensemble averaged inter-residue distances (lower triangles of Fig 3B) within conformational ensembles of Httex1. These distance maps show that for both 25Q and 46Q constructs the N17 and polyQ regions are part of a compact globule whereas the C-terminal region adopts extended structures, again consistent with the ΔCα–ΔCβ profile for 25Q. To better interpret the pattern of inter-residue distances we normalized each pair of inter-residue distances by the values obtained from simulations of Httex1 as expanded, ideal random coils. If the average inter-residue distances were lower than that of an ideal random coil, then the normalized values are less than one. Conversely, if the average inter-residue distances were equal to or larger than the corresponding distances within an ideal random coil, then the normalized values would be equal to or greater than one. These normalized inter-residue distances are shown in the upper triangular parts of Fig 3B for Q25 and Q46, respectively. The pattern of inter-residue distances is consistent with the designation of an average “*tadpole-like*” topology for Httex1 that is characterized by a compact head composed of N17 adsorbed on the polyQ tract (51) and an extended tail composed of the C-terminal region. The preference for the N-terminal and polyQ tracts to interact with one another and adopt compact conformations is consistent with the rigidity observed experimentally within this region. It is also consistent with the so-called “domain cross-talk model” that has been proposed by Kokona et al. (52) and used by Shen et al. (53) to interpret the aggregation landscapes encoded by different Httex1 constructs. Importantly, our secondary structure analysis and analysis of distance maps suggest there is limited persistent secondary structure within the compact domain encompassing N17 and the polyQ tract and these observations are consistent with the HDX data.

**Figure 3.**
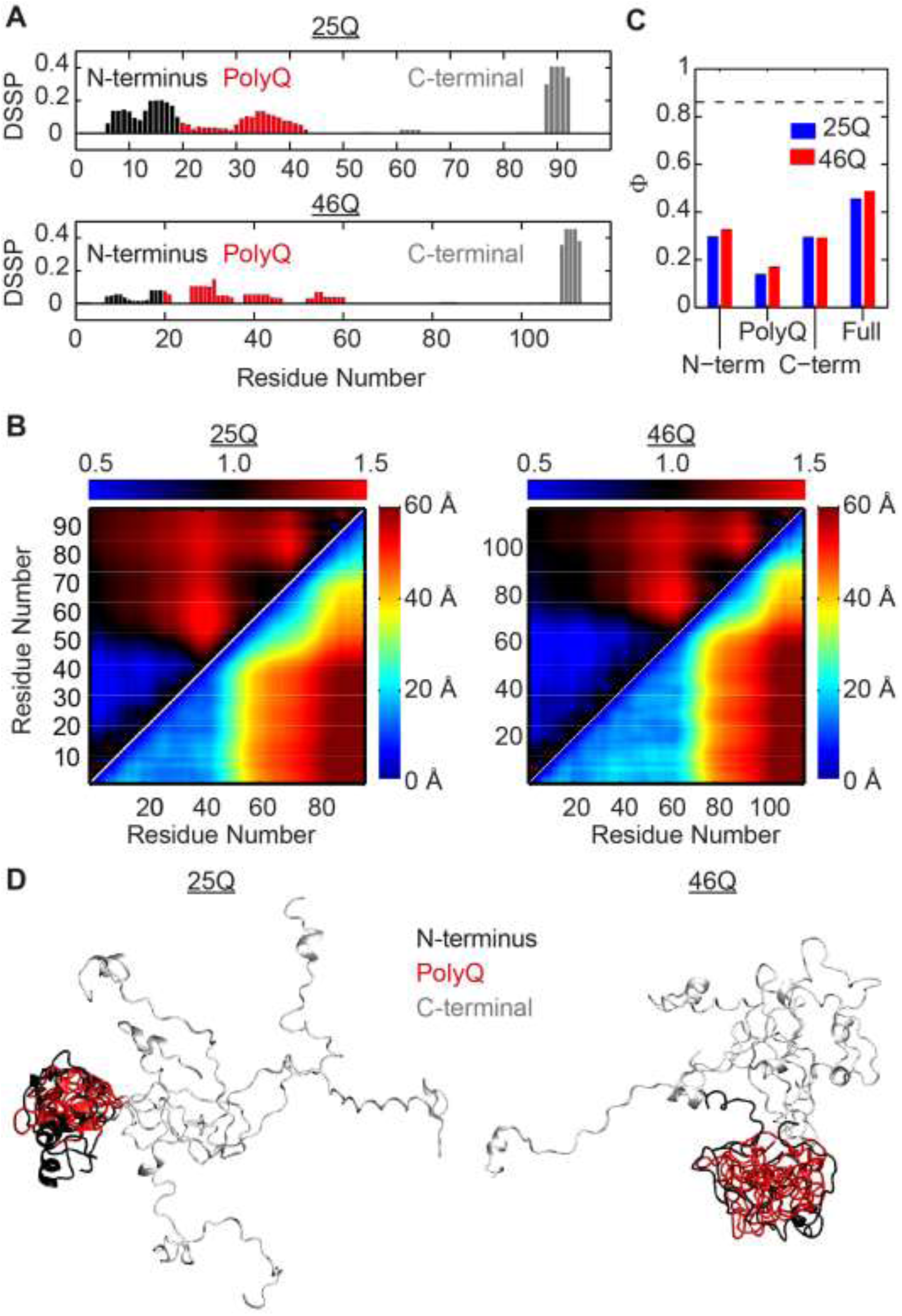
Atomistic simulation results are consistent with experimental results and suggest Httex1 adopts tadpole-like architectures. (**A**) Fraction of frames in which a residue is found in a helix of length ≥3 residues (positive) or an extended stretch of ≥3 residues (negative) as defined by the DSSP algorithm for Httex1 of 25Q and 46Q, respectively. Domains are coloured coded as shown. (**B**) The lower triangles show the average distance (in Å) between each pair of residues within the Httex1 25Q and 46Q, respectively. The hotter the colors the farther two residues are from each other. The upper triangles show the distances normalized by a reference Flory random coil for Httex1 25Q and 46Q, respectively. Here, blue colors indicate the chain is more collapsed than the reference coil and red colors indicate the chain is more expanded than the reference coil. (**C**) The degree of similarity between simulated ensembles, Φ, calculated over fragments of Httex1 (N-terminus, polyQ domain, and C-terminus) or over the full protein. Low Φ values correspond to high degrees of heterogeneity within the simulated ensembles. The dashed line corresponds to the Φ-value associated with a reference folded protein (Ntl9). (**D**) 5 representative snapshots, aligned over the polyQ domain and the first 10 residues of the C-terminus for Httex1 25Q and 46Q, respectively. Domains are coloured coded as shown.

To quantify the degree of conformational heterogeneity within Httex1, we examined the degree of conformational similarity within the entire ensemble of simulated conformations. Similarity is quantified in terms of an order parameter denoted as Φ where 0 ≤ Φ ≤ 1 (54). If the degree of conformational heterogeneity within a chain is similar to that of a reference random coil, then the extent of conformational dissimilarity within the ensemble is equivalent to that of the coil ensemble and Φ≈0. Conversely, if a chain adopts a singular, stable structure, then the conformational heterogeneity is low and Φ≈1. Fig 3C shows the Φ values computed for different Httex1 regions. The value of Φ is less than 0.4 for the N17 and C-terminal proline-rich regions for both 25Q and 46Q constructs. Additionally, the value of Φ is less than 0.2 for the polyQ region. These numbers point to significant conformational heterogeneity within the polyQ tract. Interestingly, despite significant conformational heterogeneity within each sequence module of Httex1, there is a consistent overall tadpole-like topology for the entire Httex1 sequence. This translates to higher values of Φ (≈0.5) when this parameter is calculated over the entire sequence. Importantly, the values of Φ are similar for the 25Q and 46Q constructs for all regions and the full length Httex1. The calculated sequence-specific preference for conformational heterogeneity is consistent with the chemical shifts observed for 25Q, as well as the HDX results for 25Q and 46Q. Furthermore, our results show that sequences with compact domains and a persistent preference for tadpole-like architectures are also compatible with significant conformational heterogeneity.

Overall, our structural studies uncover a tadpole-like topology of Httex1 that is encoded by distinct blocks of repetitive amino-acid sequence blocks, *i.e*., the polyQ tract versus pro-rich C-terminal domain. Representative structures from the tadpole-like ensemble are shown in Fig 3D. The overall tadpole-like architecture is preserved, as revealed by atomistic simulations across polyQ lengths. The radii of gyration and surface areas of polyQ globules increase with chain length as *N*^1/3^ and *N*^2/3^, respectively (55). Here, *N* is the number of residues within the polyQ tract. Accordingly, with increased polyQ lengths, the globular polyQ tract becomes more prominent within the tadpole-like conformations of Httex1. These results have been confirmed in recent studies that combine single molecule Förster resonance energy transfer experiments and ABSINTH-based atomistic simulations of Httex1 for five different polyQ tracts that span the spectrum of wild type and pathological lengths (56). Therefore, we propose, in accord with previous suggestions (56), that the increased prominence of the globular polyQ tract could lead to a gain of new intracellular protein-protein interactions involving the polyQ domain and low complexity domains from other proteins. We tested this hypothesis through proteomic analysis of the gain-of-function interactions engendered by expanded polyQ tracts.

### PolyQ expansion leads to gain of interactions of soluble Httex1 with the proteome

To probe how polyQ expansion modulates protein-protein interactions in the cellular context, we transiently expressed GFP-tagged Httex1 in mouse neuroblastoma (Neuro2a) cells and immunoprecipitated the soluble pool of Httex1 (which may include monomers and oligomers) for proteomic analysis of binding partners after depleting large aggregates by pelleting. Using a label-free quantification strategy, we found five proteins to be enriched by more than two-fold (p<0.05) as binding partners of 46Q when compared to the 25Q counterpart (Fig 4; Table S3). These proteins were Fus, Pebp1, Prdx6, Gars, and Hist1h4a. None of these proteins are known to interact with each other as assessed by protein-protein interactions with the STRING v10 algorithm (57). This suggested that the interactions between Httex1 and each of Fus, Pebp1, Prdx6, Gars, and Hist1h4a are independent of one another. One explanation is that they are mediated by novel properties arising from the expanded polyQ tract in 46Q.

**Figure 4.**
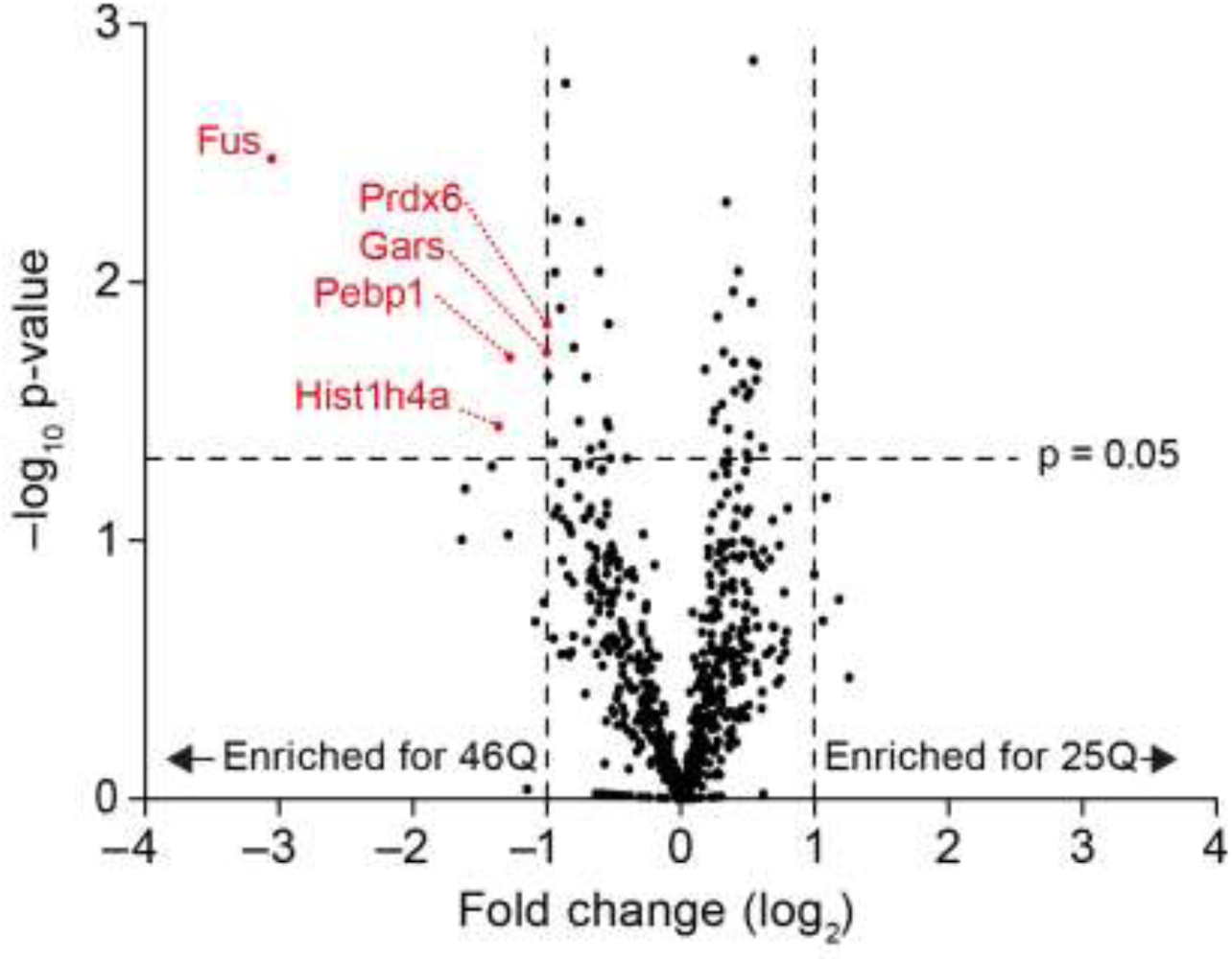
PolyQ expansion alters binding partners to soluble Httex1 states in cells. Shown is a Volcano plot of binding partners to soluble Httex1 measured from proteomic analysis of GFP-trap immunoprecipitates from Neuro2a cells transfected with Httex1-GFP. Data was acquired using label free MS/MS methods from n=3 replicates.

The most prominent interaction partner of soluble 46Q Httex1 was Fus, which functions to sequester RNA into stress granules (58, 59). The low complexity sequence regions within Fus are enriched in polar amino acids, which mediate phase separation that drives Fus into stress granules. The N-terminal low complexity sequence in particular is able to drive liquid-liquid phase separation in the micromolar range and fibril formation that leads to hydrogels in the millimolar range (60). Our data raised the prospect that soluble Httex1 has a selective affinity to Fus through a phase-separation and co-mixing mechanism between its low complexity domains and the polyQ sequence. Indeed, previous studies showed that proteins with low complexity domains, such as Fus, are preferentially recruited into aggregates formed by Httex1 with expanded polyQ tracts (61).

We performed survival curve analyses in order to determine whether the gain of function interaction between Fus and the 46Q version of Httex1 is protective or toxic to cells. Compared to a negative control GFP protein (non-fluorescent GFP Y66L derivative GFP^inv^) (62), overexpression of Fus provided a beneficial effect on survival of cells co-expressing 46Q Httex1-Cherry, suggesting that it can repress toxicity associated with expanded polyQ tracts in Httex1, providing it is overexpressed (Fig 5A). These results are in accord with the study of Ripaud et al. who showed that overexpression of proteins with Q-rich regions reduces the toxicity associated with expanded polyQ tracts of Httex1 in yeast (63).

**Figure 5.**
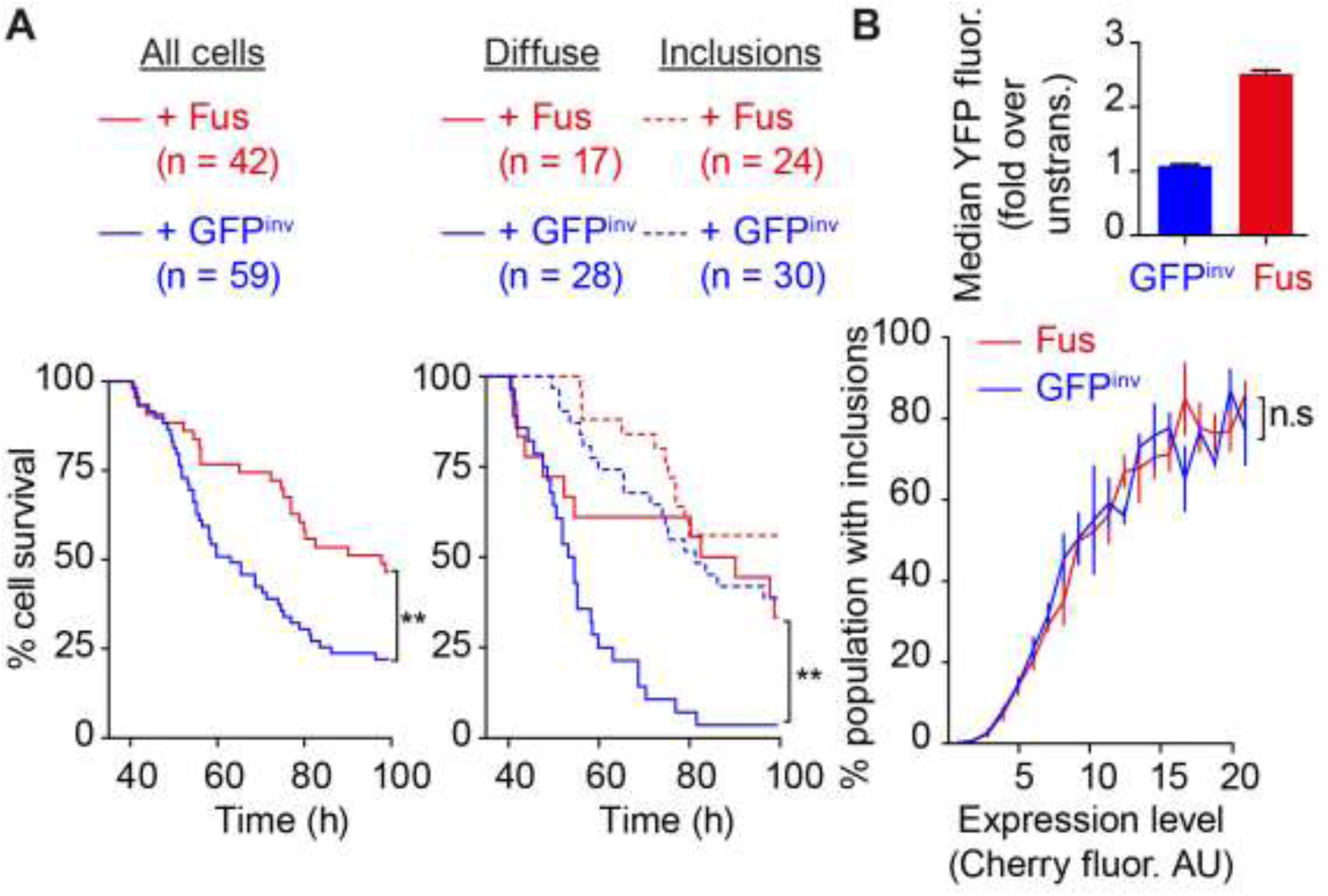
Fus suppresses toxicity mediated by soluble Httex1 but does not alter aggregation of Httex1. (**A**) Survival curves of Neuro2a cells co-transfected with Httex1 (46Q)-EmGFP and YFP-Fus or a non-fluorescent but folded GFP derivative GFP^inv^ (Y66L GFP mutant). Shown are Kaplan Meier curves for all cells (left) and for cells categorized into those that form inclusions and those that don’t (right). Mantel-Cox Log Rank test results are indicated by **, p<0.01. (**B**) Flow cytometry analysis for effect of YFP-Fus overexpression on inclusion formation by Httex1(46Q)-mCherry in neuro2a cells using pulse shape analysis (83). Shown is a plot of expression levels of Fus versus control (upper graph). Shown are percentages of cells that have inclusions as a function of Httex1 levels (lower graph; n=3, mean ± SD shown).

Next we examined whether overexpression of Fus preferentially influenced toxicity in cells with inclusions relative to cells that lack inclusions. To answer this question, we reanalysed our survival data to separate cells with inclusions from those that do not. These analyses indicated cells with soluble Httex1 died more rapidly than cells with inclusions, consistent with prior findings that soluble Httex1 generates an acute form of toxicity (7, 13) (Fig 5A). Overexpression of Fus had a substantial effect on improving the survival rate of cells with soluble Httex1 and less so on cells with inclusions. One possible explanation of this result is that overexpression of Fus detours Httex1 from homotypic interactions with itself and other deleterious heterotypic interactions. However, there appeared to be no impact of Fus overexpression on Httex1 aggregation compared to the control construct GFP^inv^ (Fig 5B), indicating that, upon overexpression, Fus becomes the dominant interaction partner for soluble Httex1. These pairwise interactions with overexpressed proteins appear to be protective, whereas gain-of-function interactions with a network of proteins might induce toxicity because of the increased interaction promiscuity through the expanded polyQ tract within Httex1.

Our minimalist hypothesis, supported by our cellular data, is that the increased prominence of the globular polyQ domain in Httex1 with longer polyQ tracts serves as an attractant for proteins with low complexity polar tracts within the cellular milieu. These gain-of-function heterotypic interactions lead to a sequestration of other proteins and Httex1 in a range of complexes. In the case of Fus, the gain of function heterotypic interactions prove to be protective upon overexpression of Fus and this leads to increased cell survival that results from depleting the soluble pool of Httex1.

## DISCUSSION

A series of prior studies established that soluble forms of polyQ expanded Httex1 are toxic to cells as quantified in cell survival measurements (7-13). Here, we have focused on a detailed structural characterization, using a combination of biophysical methods, to uncover the structural features of soluble Httex1 for wild type and pathological polyQ lengths. These studies were designed to investigate the possibility that polyQ expanded Httex1 undergoes a discernible structural change when compared to constructs with wild type polyQ. By measuring HDX by both NMR and MS we find no evidence for any changes in solvent protection, to imply the presence of stable hydrogen bonds, with increased polyQ length. However, despite the lack of persistent structure, our analysis of ^15^N{^1^H} NOEs reveals that a large fraction of the N17 and polyQ regions form conformationally rigid regions, considerably more so than archetypal intrinsically disordered proteins such as α-synuclein. These data point to the presence of compact regions within Httex1 that prevail despite the absence of specific and persistent hydrogen bonding. The apparent duality of conformational rigidity and heterogeneity are reconciled by atomistic simulations, which reveal that Httex1 adopts tadpole-like topologies regardless of polyQ length. In these tadpole-like structures, the globular head is defined by the adsorption of N17 on a globular, albeit disordered polyQ domain whereas the C-terminal proline-rich region adopts semi-flexible, rod-like conformations. The tadpole-like structures explain the impact of polyQ expansions because these expansions increase the prominence of the globular head. Specifically, the surface area and radius of gyration of the globular polyQ region increase as *N*^2/3^ and *N*^1/3^ with polyQ length *N*. This points to the prospect of gain-of-function interactions through the polyQ domain that include new heterotypic interactions that are unavailable to Httex1 with wild type polyQ and an increased tendency to aggregate via homotypic associations, which would also be weak for Httex1 with wild type polyQ.

Proteomics analysis reveals gain-of-function interactions whereby certain protein interaction partners interact more prominently with polyQ expanded Httex1 when compared to wild type Httex1. These interaction partners share the features of being RNA binding proteins that include low complexity domains, but have nothing else in common. It appears that the five proteins *viz*., Fus, Pebp1, Prdx6, Gars, and Hist1h4a primarily target the polyQ tract given that the gain of these interactions requires the presence of an expanded polyQ tract within Httex1. Of the five proteins that we identified as gain-of-function interactions, Fus was the most enriched. This protein is part of RNA stress granules and mutations within Fus are known to be associated with Amyotrophic Lateral Sclerosis (ALS) (59, 64). Previous studies showed that the low complexity disordered domain of Fus leads to preferential interactions and recruitment of Fus into aggregates with polyQ expansions (61). Here, we find that overexpression of Fus leads to increased cell survival, implying a diminution of cellular toxicity, of cells expressing Httex1 with expanded polyQ. The benefit appeared selective to cells with soluble Httex1 as opposed to cells with inclusions of Httex1. It appears that overexpressed Fus is capable of imparting protective effects on cells by preventing polyQ expanded Httex1 from engaging in promiscuous heterotypic interactions that lead to the toxicity. Through linkage effects (65), we expect that heterotypic interactions between Httex1 and overexpressed Fus modulates the intracellular aggregation and phase behaviour of Httex1 while also influencing any toxic gain-of-function heterotypic interactions. Clearly, what is required is a systematic assessment of the interplay between homotypic interactions that drive aggregation and phase separation of Httex1 (35) and gain-of-function heterotypic interactions with polyQ expanded Httex1 that could either be protective or toxic to cells. This requires assays to characterize the impact of heterotypic interactions on Httex1 phase behaviour and a framework to understand the interplay among multiple gain-of-function heterotypic interactions.

Overall, our findings converge upon a detailed structural characterization of monomeric Httex1 for wild type and pathological polyQ lengths. Our findings are in complete agreement with the recent results of Warner et al. (56) Additionally, and importantly, we have gone beyond the work of Warner et al. to show that the gain-of-function interactions can be attributed to the tadpole-like architecture of Httex1, which is characterized by a globular polyQ domain – with an adsorbed N17 – whose surface area increases monotonically with polyQ length. The gain-of-function interactions can be rendered protective through overexpression of one of the nodes in the network, as shown here via the overexpression of Fus. In general, however, the gain-of-function interactions are likely to be toxic if they sequester essential proteins in non-physiological programs, especially since the proteins that feature prominently in the gain-of-function network are transcriptional regulators that bind RNA. A detailed investigation of the multiple routes to cellular toxicity through the soluble forms of Httex1 and the design of inhibitors of toxicity is now feasible given our structural characterization that goes beyond almost all previous efforts that were limited to artificial constructs (66).

## MATERIALS AND METHODS

### Plasmids

For bacterial expression, Httex1 was cloned as a fusion with a TEV protease cleavable his-tagged maltose binding protein (MBP) tag using a pET28 vector (Novagen, Hornsby Australia) using PCR-mediated cloning procedures similar to described (62). Here, the construct lacked the fluorescent protein tag and had an appended unique tryptophan at the C-terminus (details of sequence in Table S1).

For mammalian expression, cDNA (except Fus) was expressed in pTREX-based vectors (Invitrogen) and encoded Emerald derivative of GFP (EmGFP), Httex1 25Q-EmGFP, 46Q-EmGFP, GFP Y66L, Httex1 25Q-mCherry or Httex1 46Q-mCherry, and mCherry in the pEGFP C1 vector (described in detail in Table S1) (8). Human Fus was cloned into pEYFP-C1 vector.

### Recombinant Httex1 production

To produce unlabelled or ^15^N-labelled Httex1, pET28-Httex1 plasmid was transformed into BL21 (DE3; NEB, Ipswich MA) cells using kanamycin as the selection antibiotic. A single colony was picked to create an overnight culture in 10 mL LB at 37 °C in a shaking incubator. 2 mL overnight culture was used to inoculate 500 mL autoinduction media ZM-5052 (67) containing kanamycin (50 μg/mL; Sigma-Aldrich). ^15^N-labelling was achieved using N-5052 media (67) using ^15^NH_4_-Cl (Sigma-Aldrich). Cells were grown to an OD_600_ ≈ 0.4 at 37 °C in a shaking incubator. The cells were transferred to a 16 °C shaking incubator and incubated a further 35 hours for unlabelled preps and 48 hours for ^15^N-labelled preps. Cells were harvested by centrifugation (5,000 *g*; 30 minutes; 4 °C), resuspended in Tris-HCl (100 mM; pH 7.5) with cOmplete protease inhibitor cocktail tablets (Roche) and lysed using a final concentration of 2.5 mg/mL hen egg white lysozyme (Sigma-Aldrich; stock 50 mg/mL lysozyme, 350 mM Tris-HCl, 50% v/v glycerol; pH 7.5) with vigorous shaking and stored at –20 °C for at least 16 hours.

^15^N-, ^13^C-labelled Httex1 was produced as previously described (68). ^15^NH_4_Cl and D-[^13^C] glucose (Sigma-Aldrich) were used as the nitrogen and carbon sources, respectively. Briefly, a 10 mL starter culture was prepared in LB as described above. Modified M9 media (68) (100 mL) supplemented with ^14^NH_4_Cl (50 mM), D-[^12^C] glucose (2 g), and kanamycin (50 μg/mL) inoculated with the starter culture, and this was cultured overnight at 37°C in a shaking incubator. Cells from 50 mL overnight culture were pelleted (2,000 *g*) and resuspended in modified M9 media containing ^15^NH_4_Cl (50 mM), D-[^13^C] glucose (0.3% w/v), and kanamycin (50 μg/mL). Cells were incubated in a 37 °C shaking incubator until the OD_600_ reached 0.7–0.8. Cells were induced with IPTG (final concentration 1 mM), and cultured in a shaking incubator at 16 °C until the OD_600_ plateaued – typically 24 hours post induction. Cells were harvested, lysed and stored as described above.

Proteins were purified first through a 5 mL HisTrap column (GE Healthcare, Australia) using binding buffer (20 mM Tris-HCl, 500 mM NaCl, 5 mM imidazole; pH 7.5) and elution buffer (20 mM Tris-HCl, 150 mM NaCl, 150 mM imidazole; pH 7.5) then a 5 mL MBPTrap column (GE Healthcare; on the GE Healthcare ÄKTA Pure chromatography system).

### TEV protease production, purification and cleavage

TEV protease was produced as described (69). Briefly, BL21 *E. coli* were transformed with pTEV (S219V) and grown in 2×YT broth with kanamycin as the selection antibiotic. 5 mL of overnight culture was used to inoculate 1 L 2 × YT broth, and the culture was incubated (with shaking) at 37 °C until OD_600_ ∼0.6. IPTG (0.4 mM final concentration) was added for a 4 hour induction at 30 °C. Cells were pelleted (5,000 *g*; 30 min; 4 °C) and lysed with lysozyme as described above. TEV protease was purified using a HisTrap column (GE Healthcare) using bind buffer (40 mM sodium phosphate, 300 mM NaCl, 25 mM imidazole; pH 6.8) and elution buffer (40 mM sodium phosphate, 150 mM NaCl, 300 mM imidazole; pH 6.8), then stored in 10% glycerol. This protocol produced a final concentration of 3–4 mg/mL TEV protease. Cleavage activity was tested before use.

MBP tagged Httex1 was diluted to a concentration of 20 μM and then cleaved by addition of purified TEV protease (1:1 v/v) for 40 minutes at room temperature. Httex1, which lacks the his-tag, was purified from the other fragments MBP, uncleaved MBP-Httex1 and TEV protease, which all contain his tags, by filtration through a 5 mL HisTrap column equilibrated with a low salt buffer (20 mM Tris, 100 mM NaCl, 25 mM imidazole; pH 7.5). The flow through containing Httex1 was concentrated using an Amicon Ultra centrifugal filter unit (3,000 Da NMWCO cut off; Merck-Millipore, Bayswater Australia). The resultant concentrate was desalted using a PD10 column (Sephadex G-25; GE Healthcare) into sodium acetate buffer (150 mM; pH 4) and adjusted to a final protein concentration of 100 μM as measured by spectrophotometry.

### HDX-MS sample preparation

To initiate protein deuteration 5% v/v protein solution (Httex1 25Q and Httex1 46Q) was mixed with 95% deuterated buffer at either pH 7.5 (20 mM Tris-HCl, 100 mM NaCl) or pH 4 (150 mM sodium acetate buffer). H-D exchange was quenched by reducing pH to 2.5 with DCl at five different time points: 30 seconds, 1 minute, 3 minutes, 5 minutes and 10 minutes. Each reaction was snap frozen in liquid nitrogen. Proteins were applied to a 2.1 × 10 mm column 300 Å C8 reverse phase column (Agilent Technologies, Mulgrave Australia) on ice in-line to an Agilent esiTOF Mass Spectrometer. Proteins were eluted with a gradient of buffer A: 0.1% v/v formic acid; and B: 95% v/v acetonitrile, 0.1% v/v formic acid in H_2_O at a flow rate of 0.25 mL/minute, increasing the percentage of B from 5% to 80% v/v over one minute. Mass spectra were deconvoluted using Agilent Mass Hunter, and these deconvoluted masses were compared to total deuterated backbone mass (70).

### NMR data acquisition

NMR data for Httex1 25Q were acquired on a 700-MHz Bruker Avance HDIII spectrometer with a triple resonance cryoprobe. Due to the propensity of Httex1 to aggregate, data were acquired at 5 °C. 3D HNCACB, HNCOCACB, HNCO and HNCACO experiments were acquired using ^15^N and ^13^C isotopically labelled Httex1. The acquisition parameters for all HSQC involved obtaining 1,024 points for ^1^H, 128 points for ^15^N, over 16 scans, with an acquisition time of 80 minutes. Data were processed using NMRPipe (71), and data analysed using NMRFAM-SPARKY (72). Assignments were determined using the PINE server (73, 74).

Protein (Httex1 25Q) was prepared for NMR as described above. ^15^N,^1^H HSQC was acquired on protein in protonated buffer at both pH 7.5 (20 mM Tris-HCl, 100 mM NaCl) and pH 4 (150 mM sodium acetate buffer). The same buffers were prepared using ^2^H_2_O. Directly following acquisition of the protonated buffer ^15^N,^1^H HSQC, buffer was exchanged for deuterated buffer using a NAP5 column (GE Healthcare) and another ^15^N,^1^H HSQC acquired.

^15^N{^1^H}-NOE experiments were acquired in sodium acetate buffer (150 mM; pH 4) at 5 °C with a saturation pulse of 4 seconds and an additional relaxation delay of 5 seconds (75, 76). Assignments from 3D experiments were superimposed on the ^15^N{^1^H}-NOE spectra and the differences in the saturated and reference spectra compiled using Microsoft Excel. Further analyses were performed using relax (77).

The NMR chemical shift data for ^15^N, NH, ^13^Cα, ^13^Cβ and ^13^C’ of Httex1 Q25 have been deposited at the Biological Magnetic Resonance Data Bank with accession code 27161.

### Details of atomistic simulations

Simulations of Httex1 were performed using the CAMPARI simulation package (http://campari.sourceforge.net) utilizing the ABSINTH implicit solvation model and forcefield paradigm. The move set utilized includes translational, pivot, concerted rotation, and sidechain rotation moves, as well as moves that allow for the accurate and efficient sampling of proline rings. Additional details of the implicit solvation model and move sets have been published previously. Three independent simulations were conducted for each construct using parameters from the abs_3.2_opls.prm parameter set. Protein atoms, as well as neutralizing and excess Na^+^ and Cl^-^ ions, were modeled using atomistic detail. The excess NaCl concentration in the simulations was 5 mM. The specific sequences used were Ace-GHMATLEKLMKAFESLKSF-Q_n_-P_11_-QLPQPPPQAQPLLPQPQ-P_10_-GPAVAEEPLHRPKKW-Nme. Here, *n*=25 and 46, Ace is the N-terminal acetyl unit, and Nme is the C-terminal N-methyl amide. The protonated version of histidine was used in order to mimic a solution condition of pH 4. Simulations were conducted in spherical droplets with radii of 150 Å. Temperature replica exchange was used to enhance sampling and followed the temperature schedule T=[288 K, 293 K, 298 K, 305 K, 310 K, 315 K, 320 K, 325 K, 335 K, 345 K, 360 K, 375 K, 390 K, 405 K]. Simulations consisted of 6.15×10^7^ total steps of which the first 10^7^ were taken as equilibration steps. Here, a step consists of either a Metropolis Monte Carlo move or a temperature swap. Temperature swap steps were proposed every 5×10^4^ steps. Frames used for analyses were collected every 5×10^3^ steps over the last 5.15×10^7^ simulation steps such that each simulation generated 10,300 frames. These frames were furthered filtered to remove any frames in which a residue within the polyQ domain was within 8 Å of any residue C-terminal to the P_11_ stretch given that previous single molecule Förster resonance energy transfer experiments on Httex1 constructs were inconsistent with the polyQ domain interacting with the C-terminal region.

### Proteomics: Sample preparation and MS/MS

Neuro2a cells (1.2 × 10^7^) were plated on a T150 culture flask and transfected with the pT-REx Httex1-25Q-Em or Httex1-46Q-Em vectors using Lipofectamine 2000 according to the manufacturer’s instructions (Thermo Fisher Scientific, Australia). Cells were harvested 48 hours after transfection, transferred to Lo-Bind tubes (Eppendorf, North Ryde Australia) and lysed mechanically by vortexing with silica beads in ATP depleted lysis buffer (10mM Tris-HCl, 150 mM NaCl, 0.5 mM EDTA, 1 mM PMSF, 7 mM D-glucose, 1 U/mL hexokinase, 20 U/mL benzonase, 1 cOmplete protease inhibitor cocktail tablet per 10 mL buffer; pH 7.5). Lysates were centrifuged to remove protein aggregates (21,130 *g*) and matched for equivalent GFP fluorescence (BioRad plate reader, Gladesville Australia). GFP-TrapA beads (ChromoTek, Planegg-Martinsried Germany), pre-equilibrated as per the manufacturer’s instructions with dilution buffer (100 mM Tris-HCl, 150 mM NaCl 1 cOmplete protease inhibitor cocktail tablet per 10 mL; pH 8) were added to the lysate and incubated for 2 hours at 4 °C. The beads were pelleted (2,000 *g*) and washed thrice in dilution buffer as per manufacturer’s instructions. Beads were resuspended in triethylammonium bicarbonate (Sigma-Aldrich) and snap frozen in liquid nitrogen. Proteins were eluted from the GFP-TrapA beads with 4% w/v SDS, 100 mM DTT, 100 mM Tris-HCl; pH 7.5 at 65 °C for 5 minutes. Beads were washed twice with 100 mM Tris-HCl (pH 7.5) with the beads collected by centrifugation (2,000 *g*). The wash supernatants were pooled and added to the eluent. Eluted proteins were prepared for LC-MS/MS analysis using a FASP Protein Digestion Kit (Expedeon, San Diego CA) according to the manufacturer’s instructions with two additional wash steps with Urea Sample Solution (Expedeon). Samples were dried using a benchtop CentriVap (Labconco, Kansas City MO) and resuspended in 0.1% v/v formic acid.

Peptides were analysed by LC-MS/MS using the QExactive mass spectrometer (Thermo Fisher Scientific) coupled online with a RSLC nano HPLC (Ultimate 3000, Thermo Fisher Scientific). Samples were loaded on a 100 μm, 2 cm nanoviper pepmap100 trap column in 2% v/v acetonitrile, 0.1% v/v formic acid at a flow rate of 15 μL/minute. Peptides were eluted and separated at a flow rate of 300 μL/minute on RSLC nanocolumn 75 μm × 50 cm, pepmap100 C18, 3 μm 100 Å pore size (Thermo Fisher Scientific) as described (78).

### MS/MS data analysis

MS/MS data were analysed using MaxQuant version 1.2.2.5 and the UniProt mouse protein database (accessed August 2013) for peptide identification. Strict bioinformatic criteria were used for the assignment of peptide identity; including the application of a false discovery rate of 0.01 for both proteins and peptides (79, 80) and removal of species contained within a list of common contaminants (81). Data were further analysed using Perseus version 1.5.6.0 (MaxQuant) to determine significantly enriched species in label free quantification measurements of the various IP experiments. A two-fold increase in relative abundance and a *P* value < 0.05 (Student’s t-test) were used to determine those enriched peptides/proteins.

The mass spectrometry proteomics data have been deposited in PRIDE Archive proteomics data repository (82) with dataset identifier PXD006792.

### Fus overexpression

Neuro2a cells were plated in 24-well format at a density of 1×10^5^ cells/well. The next day, cells were transfected 4 μg plasmid DNA and 4 μL Lipofectamine 2000. Plasmids were mixed at 1:1 mass ratios of pTREX Httex1-mCherry versus pEYFP-Fus (or pTREX GFP Y66L). After 36 hours incubation, cell survival was tracked by longitudinal imaging with a JuLI-Stage live cell imager (NanoEntek, Seoul Korea) with images acquired every 20 minutes for a further 55 hours using the YFP and RFP filter channels as described (13). Cells were categorized into those that formed inclusions (at some point during the timecourse) and those that did not, and their respective survival rates were compared using the Log-rank (Mantel-Cox) test. Cells that undergo division and cells with expression levels that are outside two deviations of the population mean were excluded from the analyses.

## ACKNOWLEDGMENTS

We thank Oded Kleifeld from the Infection and Immunity Program, Monash Biomedicine Discovery Institute and Department of Biochemistry and Molecular Biology, Monash University for technical assistance with the proteomics work (now at Faculty of Biology, Technion-Israel Institute of Technology, Haifa 3200003). The US National Institutes of Health supported this work through grant R01NS056114 to R.V.P.

---

**Figure S1.**
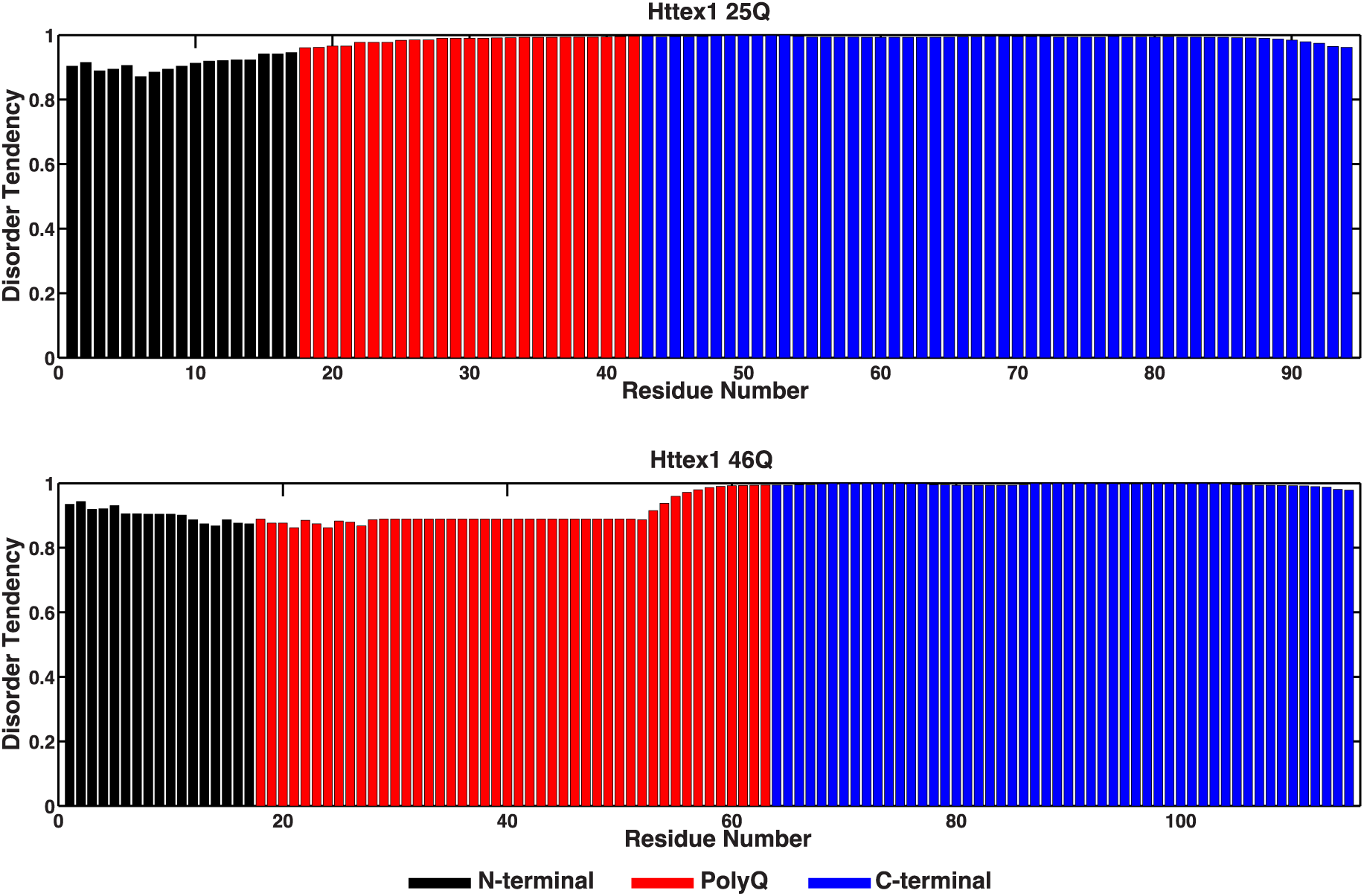
Disorder tendency predictions per residue for Httex1 25Q and 46Q as determined from the method of IUPRED (84, 85). Black bars denote N-terminal residues, red bars polyQ domain residues, and blue bars C-terminal residues.

**Figure S2.**
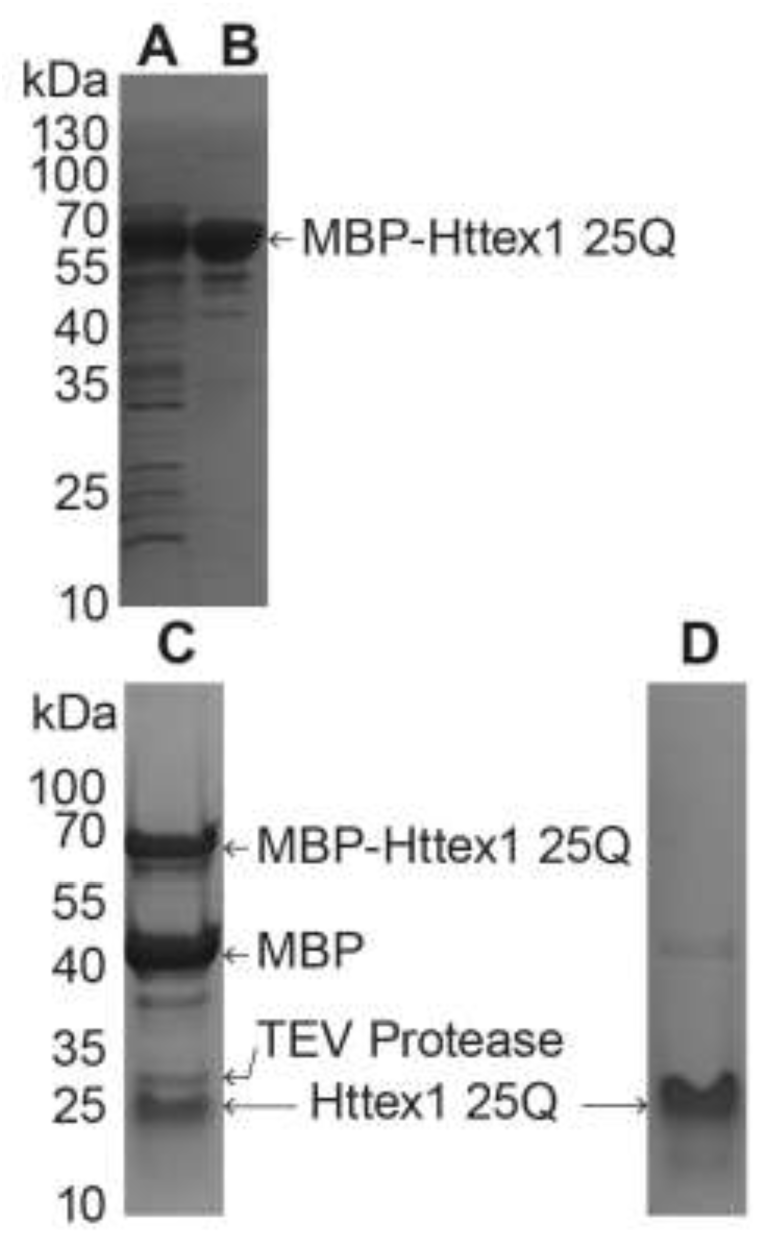
Purification of Httex1 from MBP tagged Httex1 after TEV protease treatment. Coomassie stained gels showing purification steps of Httex1 25Q. (**A**) Purification steps of MBP-Httex1 using a HisTrap column followed by (**B**) an MBPTrap column run on a Tris-Glycine 12% polyacrylamide gel. (**C**) 8-12% Tris-Tricine gel of TEV protease treated fusion protein and (**D**) after filtration of the cleavage reactants via a HisTrap column.

**Figure S3.**
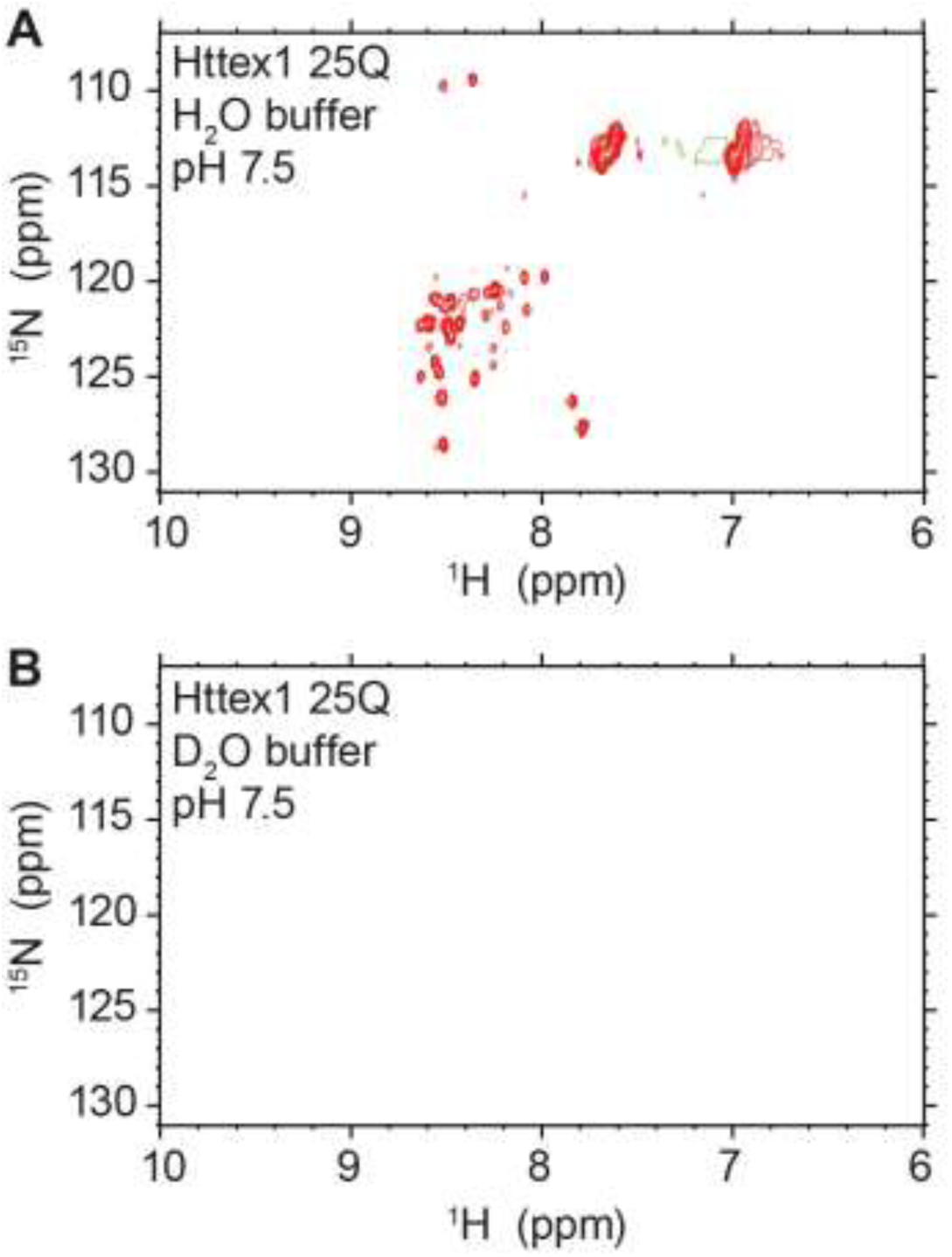
(**A**) Httex1 25Q 2D HSQC taken in low salt buffer (20 mM Tris-HCl, 100 mM NaCl, pH 7.5) at 5 °C and (**B**) in deuterated buffer.

**Figure S4.**
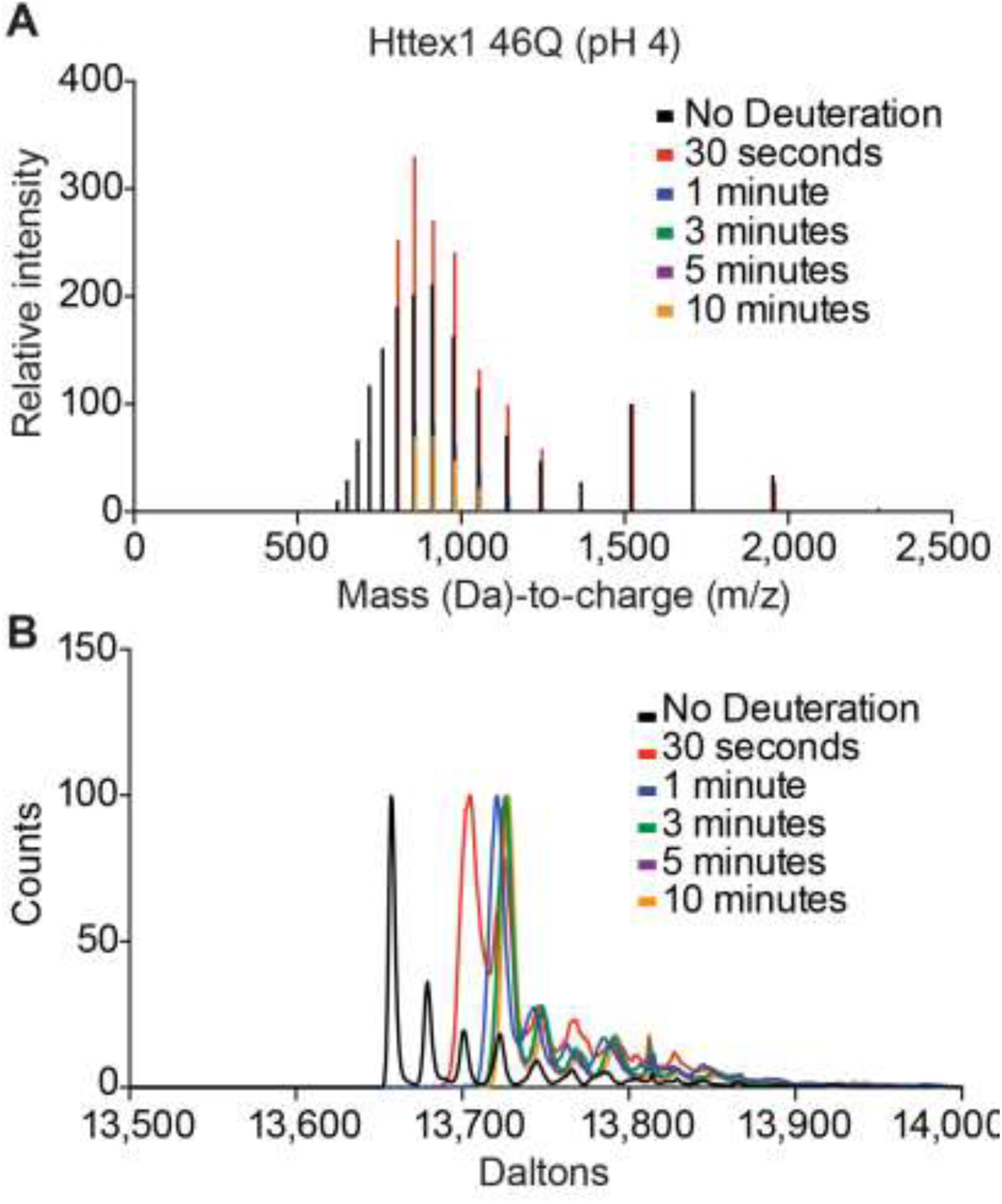
(**A**) Mass spectra of Httex1 46Q in sodium acetate buffer (150 mM, pH 4), following deuteration at different time points. (**B**) The corresponding deconvoluted mass data of the data in panel A.

**Figure S5.**
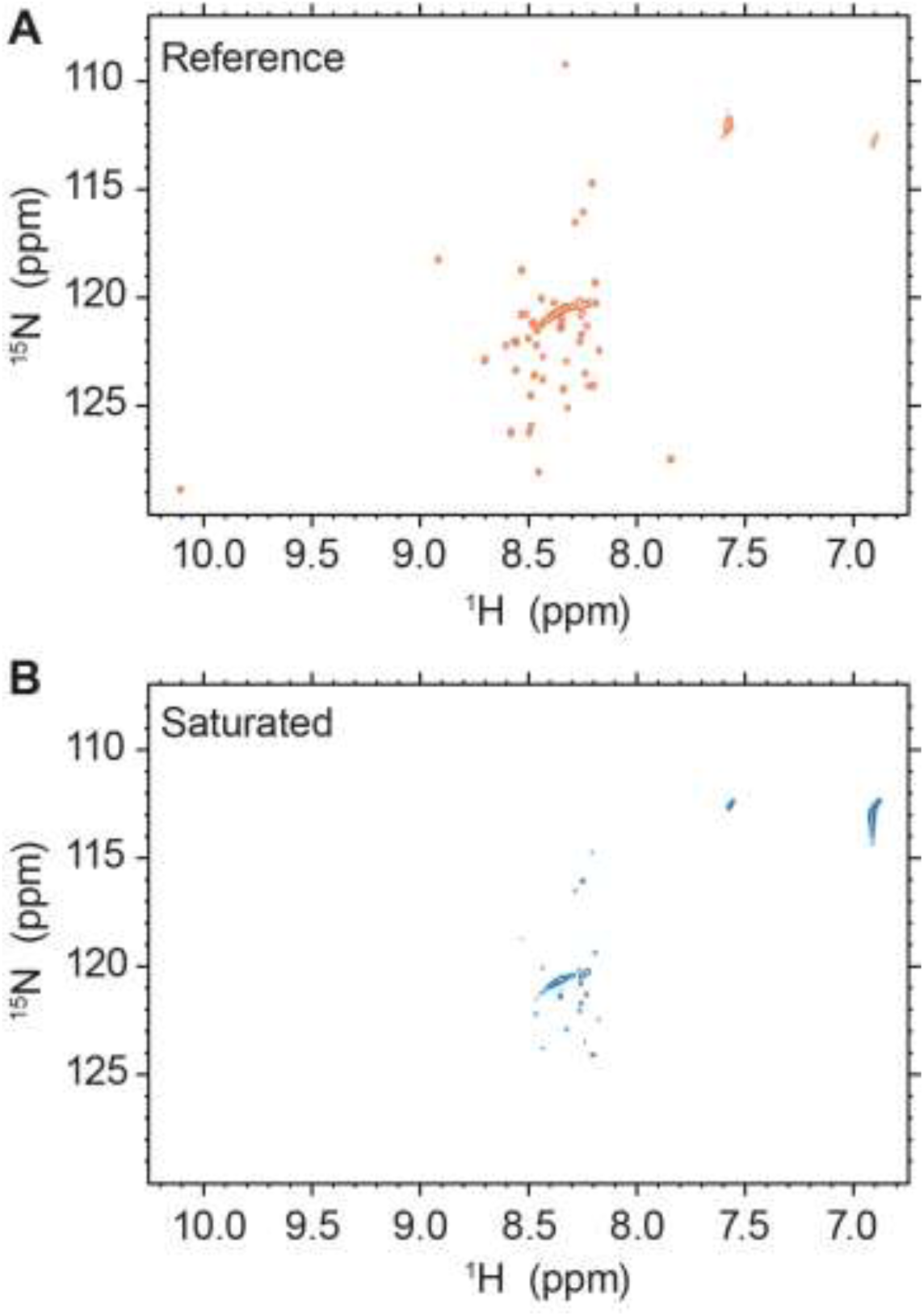
(**A**) ^15^N-^1^H-NOE reference spectra and (**B**) saturated 2D HSQC spectra for Httex1 25Q taken in sodium acetate buffer (150 mM, pH 4) at 5 °C.

**Supplementary Table 1.**
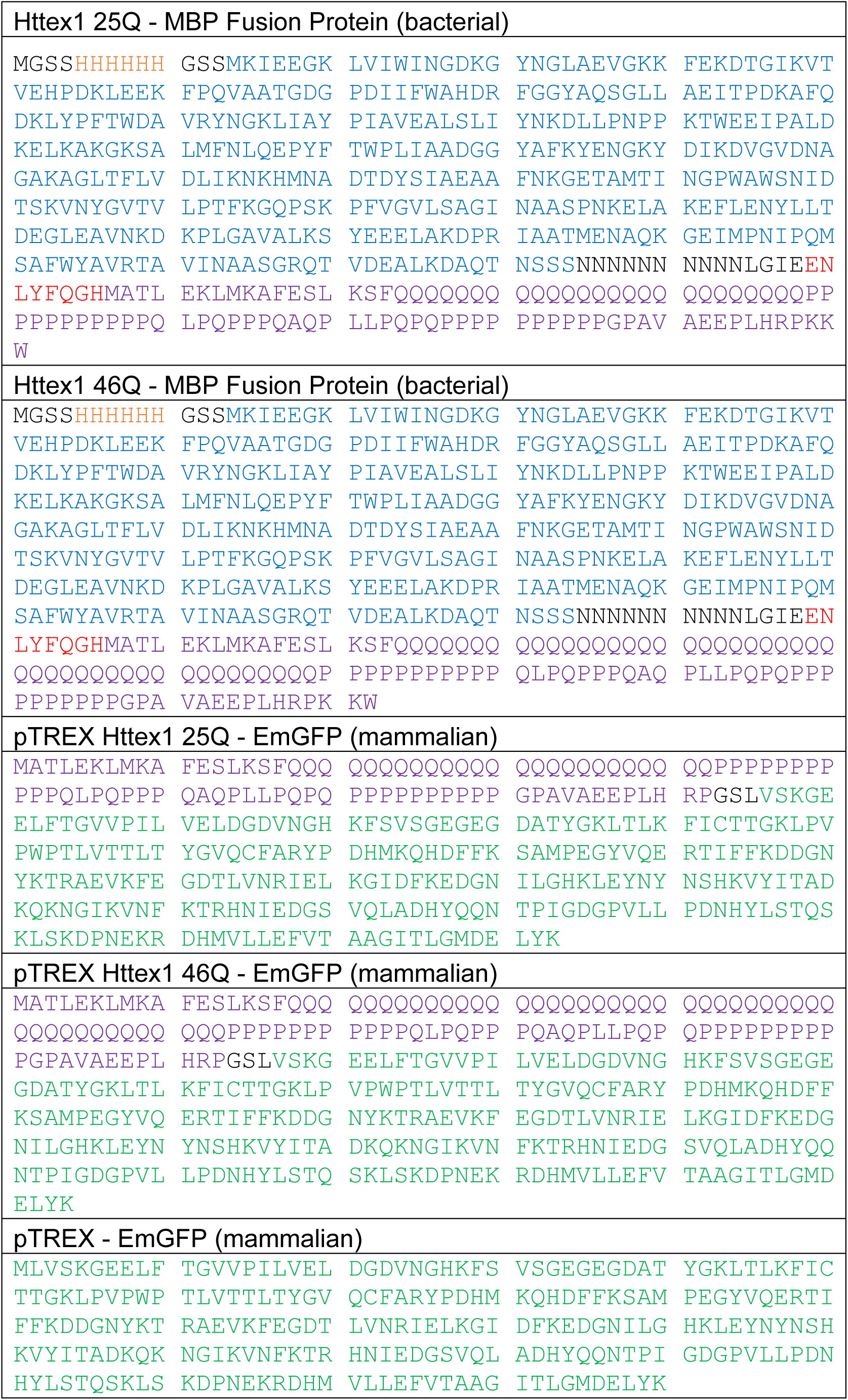
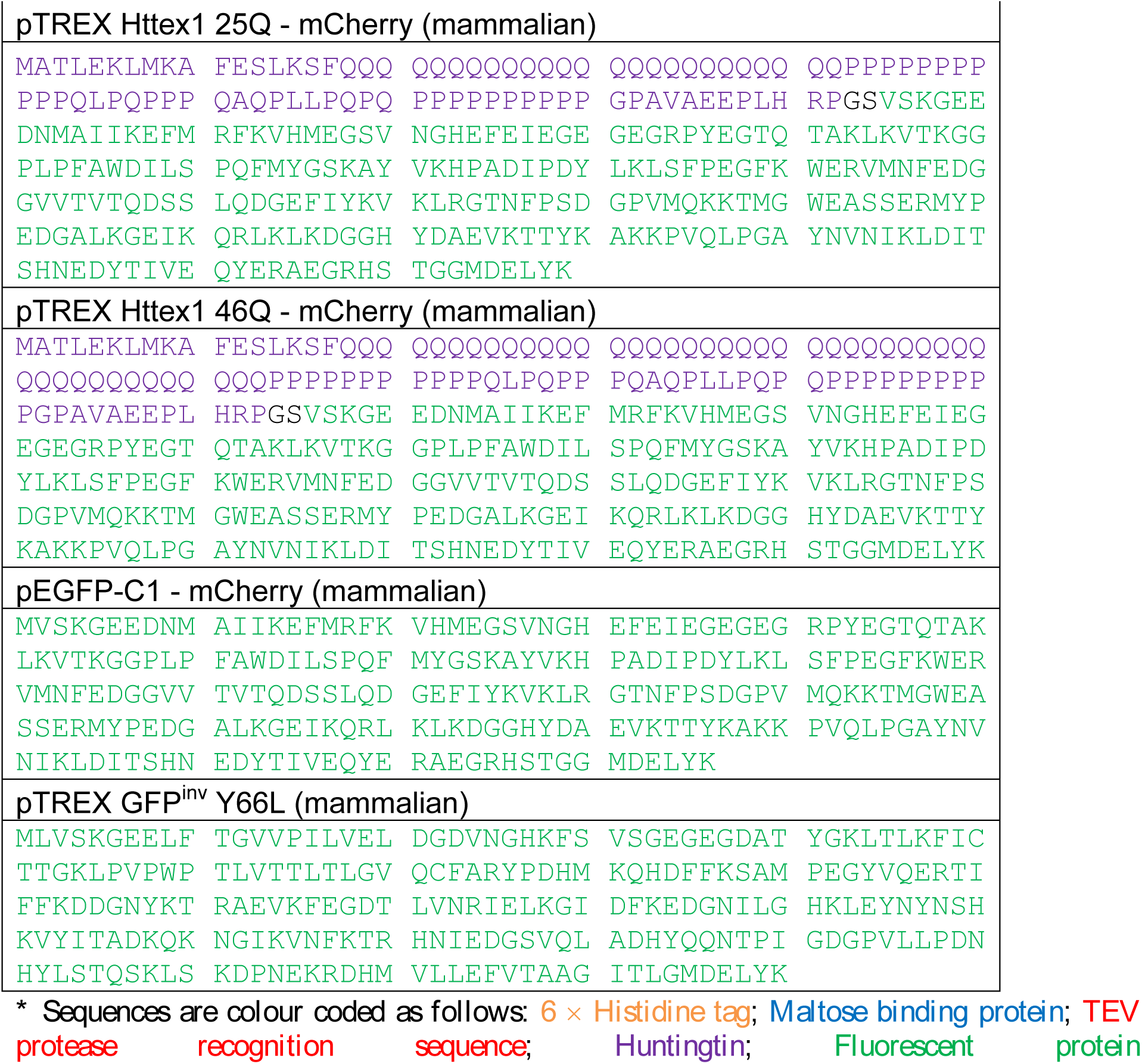
Expression vector sequences*.

**Supplementary Table 2.** Backbone assignments for Httex1 25Q.

| Assignment | <sup>15</sup>N | <sup>1</sup>H |
| --- | --- | --- |
| H02N-H | 118.252 | 8.919 |
| M03N-H | 122.897 | 8.712 |
| A04N-H | 126.244 | 8.588 |
| T05N-H | 114.797 | 8.216 |
| L06N-H | 124.234 | 8.347 |
| K08N-H | 120.566 | 8.334 |
| L09N-H | 122.063 | 8.271 |
| M10N-H | 121.39 | 8.356 |
| K11N-H | 121.753 | 8.258 |
| A12N-H | 124.075 | 8.205 |
| F13N-H | 119.426 | 8.201 |
| E14N-H | 121.325 | 8.239 |
| S15N-H | 116.563 | 8.293 |
| K17N-H | 122.483 | 8.181 |
| S18N-H | 116.082 | 8.258 |
| F19N-H | 123.01 | 8.336 |
| Q20N-H | 119.959 | 8.458 |
| Q21N-H | 120.074 | 8.249 |
| Q22N-H | 120.354 | 8.278 |
| Q23N-H | 121.43 | 8.453 |
| Q24N-H | 121.207 | 8.43 |
| QxN-H | 120.734 | 8.251 |
| QxN-H | 120.665 | 8.352 |
| QxN-H | 120.741 | 8.369 |
| QxN-H | 121.086 | 8.416 |
| QxN-H | 121.89 | 8.497 |
| Q43N-H | 120.854 | 8.385 |
| Q44N-H | 121.101 | 8.357 |
| Q56N-H | 120.341 | 8.398 |
| L57N-H | 123.383 | 8.568 |
| Q63N-H | 120.854 | 8.539 |
| A64N-H | 126.314 | 8.513 |
| L67N-H | 122.78 | 8.445 |
| L68N-H | 125.148 | 8.337 |
| Q70N-H | 122.245 | 8.62 |
| Q72N-H | 122.093 | 8.576 |
| G83N-H | 109.276 | 8.344 |
| A85N-H | 124.583 | 8.499 |
| V86N-H | 120.339 | 8.213 |
| A87N-H | 128.191 | 8.469 |
| E88N-H | 121.213 | 8.497 |
| E89N-H | 123.713 | 8.491 |
| L91N-H | 126.013 | 8.507 |
| H92N-H | 118.76 | 8.538 |
| R93N-H | 123.82 | 8.444 |
| K95N-H | 122.267 | 8.475 |
| K96N-H | 123.598 | 8.254 |
| W97N-H | 127.59 | 7.847 |
| W97-Indole | 128.853 | 10.107 |

**Supplementary Table 3.** Summary of proteomics analyses for interactors of soluble Httex1-GFP in neuro2a cells using label-free quantitation. (*Attached as a separate Microsoft Xcel file*).

